# Piping Plover Home Ranges Do Not Appear to be Impacted by Restoration of Barrier Islands and Headlands

**DOI:** 10.1101/2025.03.03.638931

**Authors:** Theodore J. Zenzal, Amanda N. Anderson, Delaina LeBlanc, Robert C. Dobbs, Geary Brock, J. Hardin Waddle

**Author notes:** Louisiana Department of Wildlife and Fisheries, Lafayette, LA. Department of Pathobiology, Wildlife Futures Program, University of Pennsylvania School of Veterinary Medicine, Kennett Square, PA.

## Abstract

Restoration of barrier island and headland habitats can alter existing and create new habitats, which may impact wildlife occupying these areas such as the threatened Piping Plover (*Charadrius melodus*). We used resight data from banded birds to develop minimum convex polygon (MCP) and kernel density estimates (KDE) of individual Piping Plover home ranges to investigate whether changes in habitat use resulted from restoration activities at Whiskey Island and Caminada Headland, Louisiana. We quantified home range areas for each season and compared changes among pre-restoration, active restoration, and post-restoration phases at each site. We had sufficient sample sizes from Whiskey Island to compare home ranges derived by MCP during the pre-restoration phase to the active restoration phase. However, we did not have enough resight data to analyze post-restoration phase by MCP or any phase by KDE at Whiskey Island. For Caminada Headland, we were able to compare all phases of restoration using MCP, but only had sufficient data to compare pre-restoration and active restoration phases using KDE. Aside from one significant decrease in core (50% isopleth) home range at Caminada Headland when comparing MCPs between post-restoration (∼8 ha) and pre-restoration (∼11 ha) phases, we found no other differences in home range size across phases at either of our study sites. The sum of all evidence generally indicating no change to Piping Plover home range size suggests that barrier island and headland restoration did not have significant positive or negative impacts. The weak response to restoration activities further suggests that birds are using similar or smaller amounts of habitat after restoration is complete and may not need to expand their foraging range following restoration. Further study can help to understand how species of conservation concern respond to coastal restoration efforts, which is critical for establishing comprehensive conservation strategies aimed at species recovery.

## Introduction

The mosaic of habitats that comprises the Gulf of America (Gulf of Mexico; *hereafter* Gulf) coast is essential to the annual cycle of over one million migrating and non-breeding shorebirds (Brush et al. 2019; Norling et al. 2012; Remsen et al. 2019; Withers 2002). The state of Louisiana contains regionally important sites for a significant number of these birds, with over 100,000 individuals documented each year (Remsen et al. 2019). Continued habitat suitability is important for supporting current levels of shorebird abundance and diversity, as many species exhibit high interannual site fidelity during the migration and stationary non-breeding seasons, and tend to concentrate in localized areas that provide abundant food resources and safe roost locations (Brush et al. 2019; Withers 2002). Diverse habitats in close proximity allow shorebirds to frequently move between habitat types within and among barrier islands, which can coincide with changing habitat availability in relation to daily and seasonal tide cycles, winds, and/or biological needs (e.g., foraging, loafing, breeding; Brush et al. 2019; Withers 2002).

The Gulf is a crucial non-breeding area for the Endangered Species Act listed Piping Plover (*Charadrius melodus*), which spends up to 70% of the annual cycle on the non-breeding grounds and exhibits high site fidelity to these locations (Drake et al. 2001; Elliott-Smith and Haig 2020; Noel and Chandler 2008). Based on data from the International Winter Piping Plover Census, 65-93% of Piping Plovers that breed within the Interior and Great Lakes regions spend the non-breeding season in the Gulf region, especially in coastal Texas and Louisiana (Brush et al. 2019; Elliott-Smith and Haig 2020). Approximately 10,000 ha of non-breeding Piping Plover Critical Habitat has been designated in Louisiana, including barrier islands that support 85% of the state’s non-breeding population (Elliott-Smith and Haig 2020; Remsen et al. 2019). The average annual count of non-breeding Piping Plovers in Louisiana was approximately 250 individuals from 1991 to 2014, with a high count of 750 in 1991 and a low of 86 in 2011 (Elliott-Smith and Haig 2020; Remsen et al. 2019). However, additional information could help to inform species recovery, such as increased understanding of habitat use during the migration and non-breeding phases of the annual cycle (Brush et al. 2019; U.S. Fish and Wildlife Service 2015).

Home range estimates can help inform shorebird conservation by identifying important sites, providing baseline data on species-habitat relationships, and evaluating impacts of management actions on shorebird behavior and ecology (e.g., Choi et al. 2014, 2019; Clemens et al. 2014; Handmaker et al. 2024; Jourdan et al. 2021). Understanding how Piping Plover home range size responds to barrier island and headland restoration is important to assess how habitat change (i.e., creation of new or increase in existing habitat types, pumping of dredged material, increase in elevation) might influence space use, given that sand placement can alter normal coastal processes and cause loss of macroinvertebrate prey (Brush et al. 2019; U.S. Fish and Wildlife Service 2015). Change in home range size may indicate a change in resource/niche availability or higher competition for use of space that may be a result of restoration activities. In Texas, Piping Plovers, which are primarily located on algal flats and lower sand flats, showed strong site fidelity, with mean core home ranges (based on 50% of location data) of 2.9 km^2^ and a linear distance moved of 3.3 ± 0.5 km (Drake et al. 2001). Studies on the Atlantic Coast determined that Piping Plover activity was concentrated in 2.2 km^2^ and 3 km^2^ of the entire study area in North Carolina and Georgia, respectively (Cohen et al. 2008; Noel and Chandler 2008). To our knowledge, there has been no analysis of Piping Plover home range within any of the central Gulf states (i.e., Louisiana, Mississippi, and Alabama).

The Louisiana shoreline is undergoing coastal protection measures, including barrier island and headland restoration, to 1) create, maintain, and restore land; 2) mitigate threats to coastal infrastructure (e.g., flooding); and 3) sustain habitats that support recreational and commercial activities (Coastal Protection and Restoration Authority of Louisiana 2023). While these restoration projects can benefit flora and fauna along the coastline, the main objective of these nearshore and offshore projects are to protect the mainland from climate change (e.g., sea-level rise; increase in tropical storm frequency and severity) and coastal land loss. Therefore, loss of barrier islands can have consequences on human life and property in addition to loss of flora and fauna. Barrier island restoration entails depositing offshore dredge material to improve island integrity, but consequently reduces benthic invertebrate abundance and alters sediment composition, partially impacting shorebird food resources and habitat availability (Gibson et al. 2018; Schulz and Leberg 2019; Wilber et al. 2009; Wilber and Clarke 2007). Restoration projects in the non-breeding area that result in engineered barrier island and beach habitats may benefit the species (Guilfoyle et al. 2006, 2007; Yozzo et al. 2004), directly addressing two of the greatest threats to Piping Plover, which are habitat degradation and sea-level rise (Brush et al. 2019; Gratto-Trevor and Abbott 2011; U.S. Fish and Wildlife Service 2015). The effects of engineered habitats on Piping Plover habitat use have been studied in the breeding region (e.g., Catlin et al. 2011, 2015; Hunt et al. 2017, 2018), but remain largely unknown in non-breeding areas (but see Bergquist et al. 2011).

Non-breeding survival of Piping Plovers is highly dependent on habitat conditions, including prey abundance, and a Piping Plover’s experience in an engineered non-breeding habitat can carry over to influence fitness during the breeding season (Gibson et al. 2018; Roche et al. 2010). Considering the amount of restoration occurring in non-breeding areas, like the northern Gulf, more information on non-breeding use of engineered habitats before, during, and after restoration may be beneficial for species recovery efforts, considering that barrier island and headland restoration activities have the capacity to negatively impact Piping Plovers (Brush et al. 2019; U.S. Fish and Wildlife Service 2015). For example, resource managers and conservation planners might use such information to design restoration projects that meet anthropogenic objectives and also provide functional habitats that benefit the Piping Plover. As engineered habitats become increasingly widespread, studies can help to evaluate if and how non-breeding habitat restoration affects Piping Plover habitat use.

Despite concerted survey efforts for Piping Plover in non-breeding areas, determining habitat use among restoration phases (i.e., before, during, and after) in the non-breeding season has not, to our knowledge, been investigated. We used resight data from banded birds to develop kernel density (KDE) and minimum convex polygon (MCP) estimates of individual Piping Plover home ranges and investigated whether spatiotemporal changes resulted from restoration activities at Whiskey Island and Caminada Headland, Louisiana. We quantified home range areas for each season (July-May) using 95% and 50% isopleths (i.e., polygons made up of 95% or 50% of resight instances, respectively) and compared changes amoung pre-restoration, active restoration, and post-restoration phases across multiple seasons. We hypothesized that home range area differs among the three restoration phases. Given previous research indicating changes to benthic invertebrate community composition and abundance following restoration, and potentially long durations between sand placement and recovery (Bergquist et al. 2011; Bolam 2011; Bolam and Rees 2003; Copertino et al. 2022; Maurer et al. 1986; Wilber et al. 2009; Wilber and Clarke 2007), we predicted that Piping Plover would increase home range area during the active and post-restoration phases when compared to pre-restoration phase as more foraging area may be necessary to meet their energetic needs.

## Methods

### Study Sites

Caminada Headland (29.1387°N, -90.1343°W) is located south and east of Port Fourchon, Louisiana in Lafourche and Jefferson parishes. The headland is ∼21 km of beach and intertidal habitat that extends from Belle Pass and continues eastward to Caminada Pass on Elmer’s Island (Figure 1). Caminada Headland is characterized by: 1) oceanside intertidal and beach habitats facing the Gulf, 2) an interior comprised of unvegetated dune and unvegetated flat habitats, and 3) bay-side habitats that include estuarine emergent marsh, mangrove, intertidal, meadow, and scrub/shrub habitats (see Table 3 in Enwright et al. 2020 for habitat definitions). From 1855 to 2015, the headland experienced an average shoreline loss of ∼12 m per year (Byrnes et al. 2018). Caminada Headland was restored through two projects that spanned from June 2013 to April 2017 (see Coastal Engineering Consultants Inc. 2017, 2015; LeBlanc et al. 2023 for more details). The barrier island restoration, performed by Weeks Marine, Inc. (Houma, LA, USA), involved bringing materials to the site via barge then resuspending the sediment and pumping it onshore (see Coastal Engineering Consultants Inc. 2015, 2017). The objective of the restoration was to preserve coastal habitats, re-establish littoral sand transport, and reduce storm impacts on Port Fourchon and Louisiana Highway 1 (Folse and Lee 2016). The restoration involved dredging ∼8.3 million m^3^ of offshore sediment to create ∼429 ha of beach and dune habitat across two project increments (LeBlanc et al. 2023). The total restoration effort increased the land area of Caminada Headland from 10.87 km^2^ in 2012, just prior to restoration, to 11.90 km^2^ in 2019 (Thurman et al. 2023).

**Figure 1.**
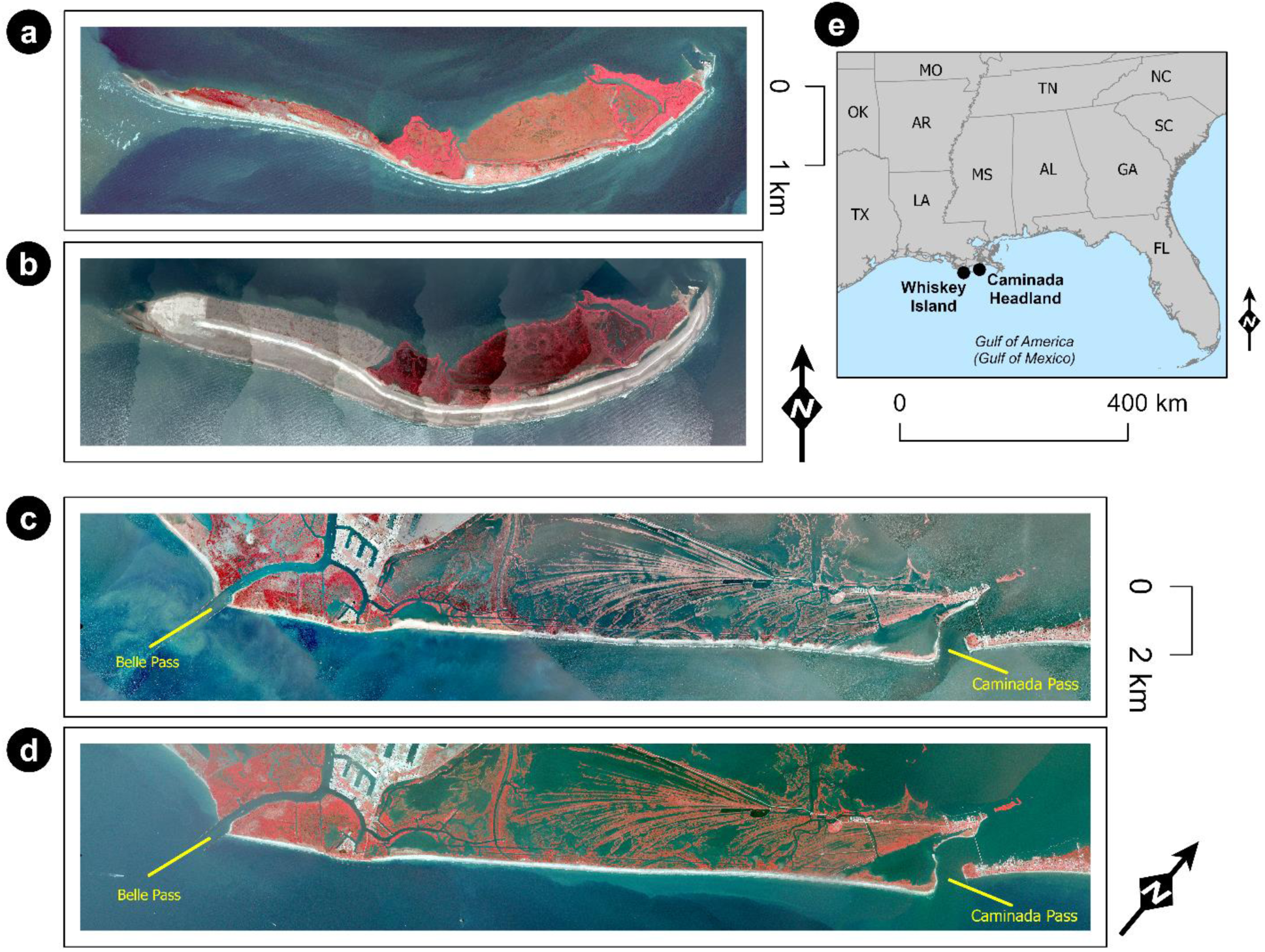
Whiskey Island prior to restoration (a; imagery from the U.S. Geological Survey [USGS], 2016) and post-restoration (b; imagery from the Louisiana Coastal Protection and Restoration Authority, 2018). Caminada Headland prior to restoration (c; imagery from the USGS, 2012) and post-restoration (d; imagery from the USGS, 2018). Location of the two sites in coastal Louisiana (e). Imagery is shown as false color, which means redder areas indicate greener vegetation.

The Isles Dernieres Barrier Islands Refuge (IDBIR), managed by the Louisiana Department of Wildlife and Fisheries, is made up of four barrier islands, which are located south of Cocodrie, Louisiana in Terrebonne Parish. Whiskey Island (Figure 1; 29.045983°N, -90.840389°W), is part of the IDBIR complex and consists of: 1) oceanside beach and intertidal habitats facing the Gulf; 2) intertidal, mangrove (*Avicennia germinans*), and estuarine emergent marsh habitats along Caillou Bay; and 3) spits of beach, intertidal, and unvegetated flat habitats extending from the east and west ends of the island. Nontidal habitats like vegetated dune, meadow, and unvegetated flats are present on the interior of Whiskey Island (see Table 3 in Enwright et al. 2020 for habitat definitions). In low-lying flats and along the edge of the mangrove and estuarine emergent marsh habitats, water from tides, rain, and other flooding events often generate tidal inlets and ephemeral pools within the island. From 1855 to 2015, ∼16 m of shoreline loss has occurred each year on average (Byrnes et al. 2018). The most recent restoration project at Whiskey Island occurred from December 2016 through May 2018 (see Coastal Engineering Consultants Inc. 2018; LeBlanc et al. 2023 for more details). The barrier island restoration, performed by Great Lakes Dredge and Dock Company, LLC (Oak Brook, IL, USA), involved pumping sediment onshore directly via submerged pipeline (Coastal Engineering Consultants Inc. 2018). Compared to the Caminada restoration project, Whiskey Island had a back barrier marsh platform component and had more sand placed in front of the dunes. The restoration involved dredging ∼8 million m^3^ of sediment from offshore sources to create 386 ha of beach, dune, and marsh habitats as well as vegetation plantings following restoration to reduce wave and tidal energy to protect the mainland shoreline from continued erosion (LeBlanc et al. 2023). This restoration effort increased the land area of Whiskey Island from 2.70 km^2^ in 2016, just prior to restoration, to 4.46 km^2^ in 2019 (Thurman et al. 2023).

At both sites, materials were being pumped onto the shoreline during most Piping Plover surveys that occurred during the active restoration periods (see Leberg and Lee 2023). Investigation of construction impacts from restoration activities on shorebirds showed little impact at our sites (Leberg and Lee 2023). While at Caminada Headland, the closest Piping Plover tended to be farther from discharge sites than random control sites, and there was no difference in distance to discharge or random sites for the three closest Piping Plovers (Leberg and Lee 2023). Leberg and Lee (2023) concluded that the inconsistency in the response of the single closest Piping Plover compared to the three closest Piping Plovers suggests most birds on the beach were not responding strongly to construction. Based on these conclusions, we assume that construction likely did not impact home range size of Piping Plover in our study.

### Field Surveys

From July through May from 2012 to 2019, avian ecologists from the Barataria-Terrebonne National Estuary Program conducted non-breeding bird surveys at Caminada Headland 2–3 times per month. All surveys occurred within two days of designated International Shorebird Survey (ISS) census dates (U.S. Fish and Wildlife Service 2015). Over the span of data collection, some surveys were missed due to logistical challenges (e.g., weather, personnel availability). Observers surveyed all suitable sparsely vegetated and non-vegetated habitats (e.g., beach, unvegetated flat, intertidal, meadow) to document several focal species including Piping Plover. Surveys were conducted from Belle Pass eastward to Caminada Pass. The study site was divided up into 4–5 sections, and each section was surveyed on foot by 1–4 observers in a single day walking the length of each section while using binoculars and spotting scopes to identify Piping Plovers. For each Piping Plover, observers recorded: 1) number of individuals; 2) leg-band combinations and alphanumeric flag codes; 3) geographic coordinates; 4) site; 5) location (i.e., bay, gulf, interior); and 6) behavior (e.g., foraging, roosting, breeding.

From 2012 to 2020, avian ecologists from the U.S. Geological Survey conducted bi-weekly surveys at Whiskey Island from August through February to monitor non-breeding bird populations, and weekly surveys from March through July to monitor breeding populations. Over the span of data collection, some surveys were missed due to logistical challenges (e.g., weather, personnel or resource availability, COVID-19 pandemic). The survey protocol included two observers independently surveying all accessible habitat (gulf-side, bay-side, interior) by foot in a single day, using binoculars and spotting scopes to document various focal species, including Piping Plovers. For each Piping Plover, observers recorded: 1) number of individuals; 2) leg-band combinations and alphanumeric flag codes; 3) geographic coordinates; 4) island zone (i.e., bay, gulf, interior); 5) habitat type (e.g., algal flat, sand flat, backshore beach); 6) behavior (e.g., foraging, roosting, breeding); and 7) substrate (e.g., moist, dry, wet, wrack, shell).

### Statistical Analysis

To assess if home range size differed among various phases of restoration, we converted repeat observations of the same individual into a generalized footprint and determined its area. Home range footprints were developed from resighted individuals using two approaches: MCP and KDE. MCP provides a simple method of estimating home ranges by creating a polygon based on the smallest area that includes a pre-specified proportion of geographic points, whereas the KDE provides a density estimate based on the clustering of geographic points. We selected two approaches because the MCP allowed additional comparisons given the lower repeated observation threshold, but the KDE provides a more detailed picture of habitat use when data are available (Gregory 2017). For MCPs, we included known individuals (i.e., high confidence of unique band combination) that had at least five observations within a single season. For KDEs, we included known individuals that had at least 19 observations within a single season (Silverman 1998).

For each approach, we characterized home ranges based on 50% and 95% isopleths. Each isopleth represents the corresponding percentage of locations where individuals are recorded, with the 50% isopleth often considered the “core area” of habitat use. To convert observed locations of birds into spatial objects, we used the “sp” (Bivand et al. 2013; Pebesma and Bivand 2005), “adehabitatHR” (Calenge 2006), “raster” (Hijmans 2022), and “rgeos” (Bivand and Rundel 2021) packages, all within the R statistical language (version 4.1.0; R Core Team 2021). To calculate the MCPs and KDEs, we used the “adehabitatHR” package (Calenge 2006) and input the desired isopleth size. The minimum sample size thresholds for calculating home range area precluded analysis of MCPs from post-restoration at Whiskey Island, KDEs for any data from Whiskey Island, and KDEs from post-restoration at Caminada Headland (see Table 1).

**Table 1.** Number of Piping Plover home ranges from each study site used in the analysis based on minimum thresholds for minimum convex polygon (MCP; n > 4) and kernel density estimation (KDE; n > 18) approaches. Some individuals were detected multiple years, thus contributing more than one home range when data were available to produce a home range for each year they were observed (see text for details).

| Site/Approach | Pre-restoration | Active restoration | Post-restoration |
| --- | --- | --- | --- |
| Whiskey<br>Island/MCP | 12 | 5 | 0 |
| Whiskey<br>Island/KDE | 0 | 0 | 0 |
| Caminada<br>Headland/MCP | 82 | 67 | 40 |
| Caminada<br>Headland/KDE | 10 | 6 | 0 |

We analyzed home range areas for the 50% and 95% isopleths using mixed effects linear models. For each possible study site and approach (50% MCP, 95% MCP, 50% KDE, and 95% KDE) combination, we selected home range area as the response variable, restoration phase as the explanatory variable, and leg band combination as a random factor to control for individuals detected across multiple years. We log transformed our response variable in each analysis to meet model assumptions. We followed each model with a Wald test to determine significance of the explanatory variable coefficients (Fahrmeir et al. 2013). If we found a significant effect of restoration phase, we used a Tukey test to determine between-group differences *post-hoc* (Tukey 1949). Since the data in our mixed effects linear model for the 50% MCP isopleth at Caminada Headland was unable to support the random effect structure, we reran these data as a linear fixed-effects model. Outputs of the linear model and mixed effects linear model were similar, hence, we present the results of the mixed model since it is more appropriate with the repeated measures approach we used. We performed analyses using the “car” (Fox and Weisberg 2019), “lme4” (Bates et al. 2015), and “multcomp” (Hothorn et al. 2008) packages. Data are available from Zenzal et al. (2025).

## Results

Sample sizes at Caminada Headland allowed for analysis of home range area across all three restoration phases with MCP (Table 1; Figures S1-S2). For KDEs, post-restoration resights were limiting, which only permitted comparisons of pre-restoration and active restoration periods (Table 1; Figures S3-S4). Restoration phase was significantly related to Piping Plover core habitat area (50% isopleths) based on MCPs (χ^2^ = 7.55; df = 2; *p* = 0.02; Figures 2a and S1). Home range area was smallest during the post-restoration phase (8.44 ± 16.54 ha; this and all following are mean ± standard deviation) and largest during the active restoration phase (14.17 ± 30.36 ha). A *post-hoc* test only determined significant core home range differences between pre-(10.77 ± 16.68 ha) and post-restoration phases. Despite seemingly large differences in overall home range area (95% isopleths) between the pre-(47.98 ± 62.87 ha), active (106.41 ± 183.01 ha), and post-restoration (31.14 ± 32.89 ha) phases, there was no statistically significant relationship between restoration phase and overall home range area, possibly due to large variances (χ^2^ = 2.88; df = 2; *p* = 0.24; Figures 2b and S2). When comparing 50% isopleths from KDEs, we found no difference between pre-(439.38 ± 723.09 ha) and active (535.25 ± 686.70 ha) restoration phases (χ^2^ = 0.26; df = 1; *p* = 0.61; Figures 3a and S3). Similarly, when comparing the 95% isopleths from KDEs, we found no difference between the pre-(2,349.24 ± 4,137.67 ha) and active (2,675.75 ± 3,051.70 ha) restoration phases (χ^2^ = 0.35; df = 1; *p* = 0.56; Figures 3b and S4).

**Figure 2.**
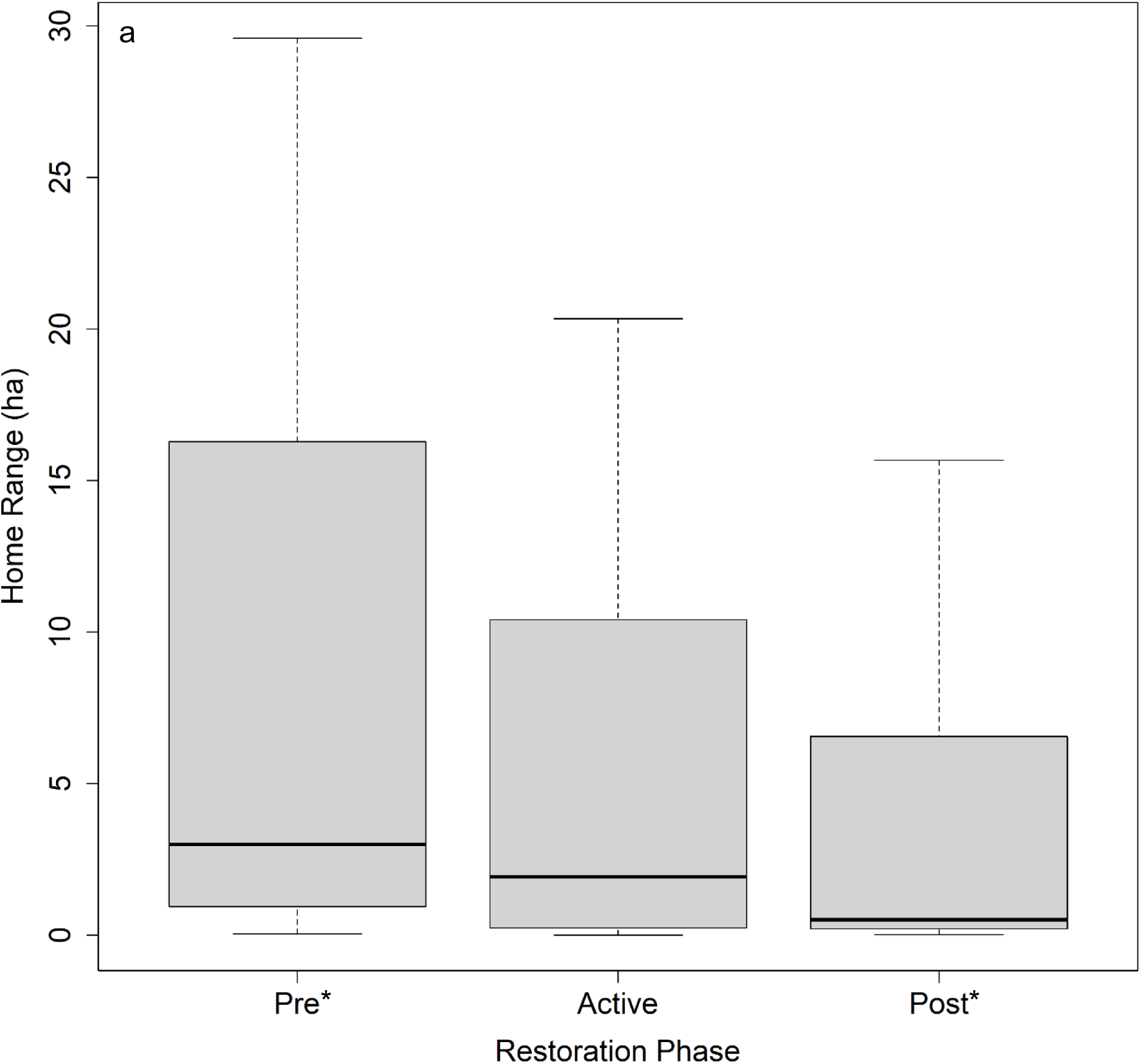
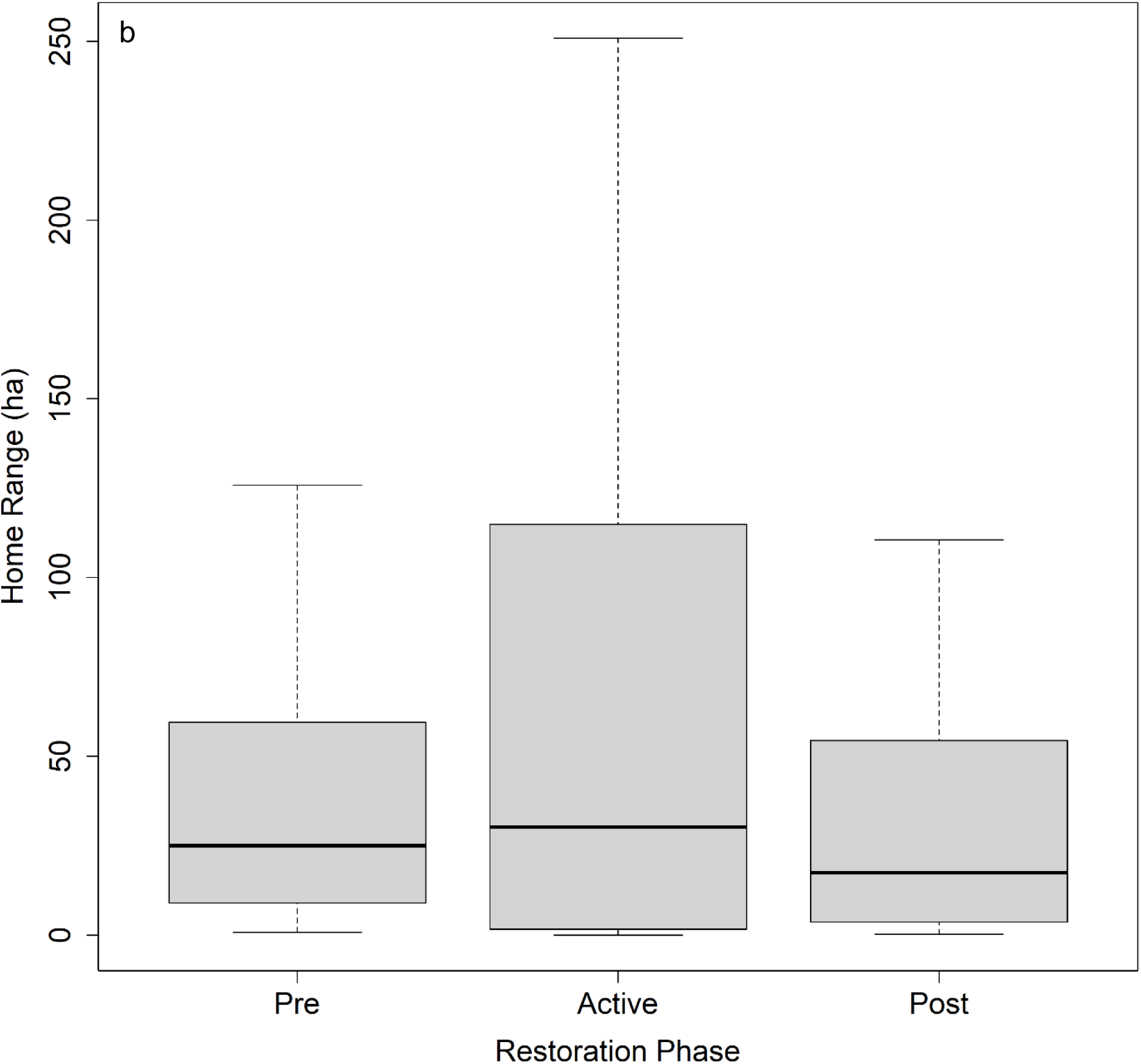
Home range size (ha) of Piping Plovers pre-restoration, during (active), and post-restoration at Caminada Headland, Louisiana based on minimum convex polygon. Displayed are the (a) 50% isopleths and (b) 95% isopleths. Central black line indicates median, top and bottom of box indicate interquartile range, and whiskers indicate total range. In the 50% isopleth analysis (a), pre-restoration and post-restoration differed significantly, which is indicated by asterisks (*); we found no other significant differences.

**Figure 3.**
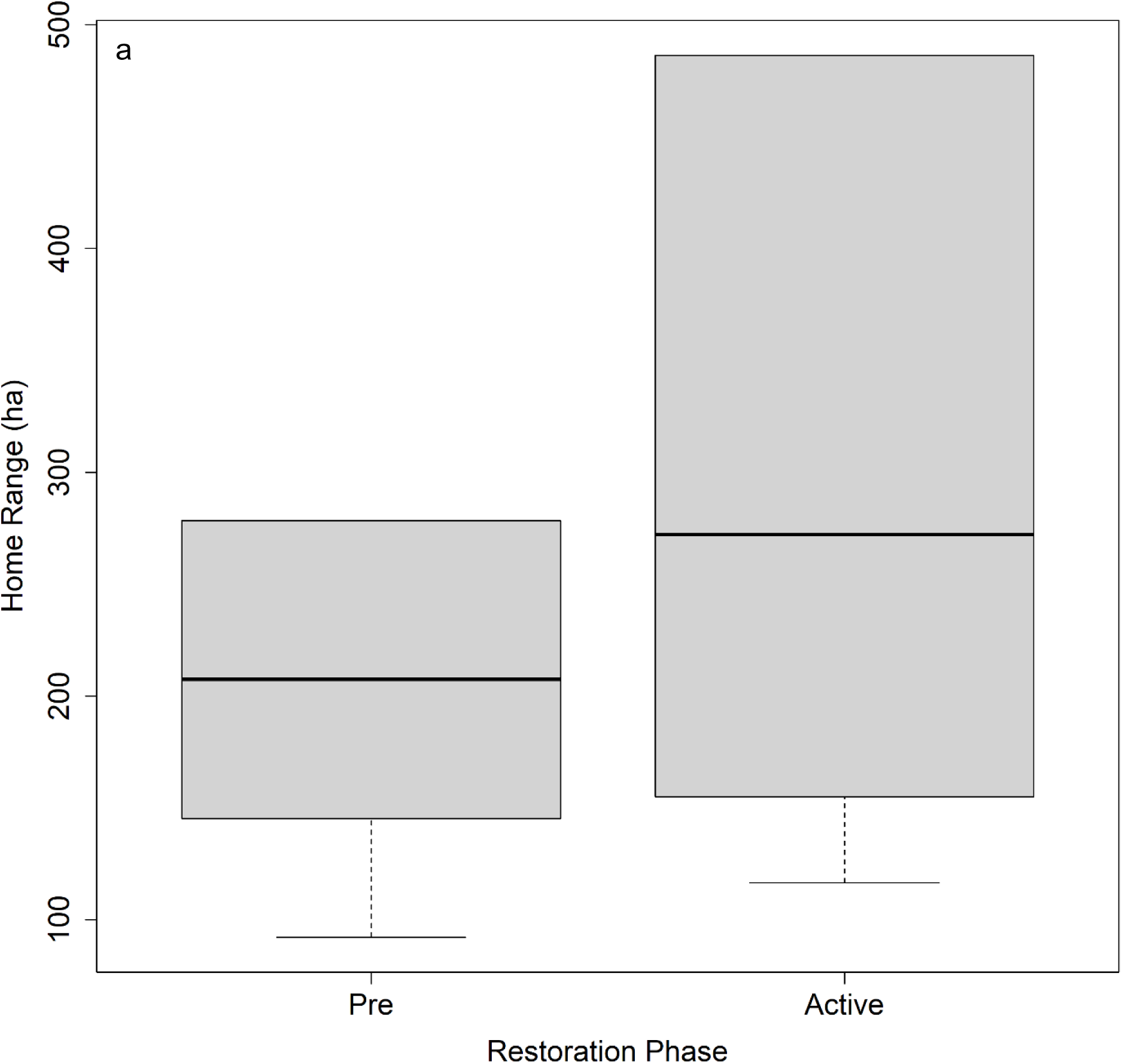
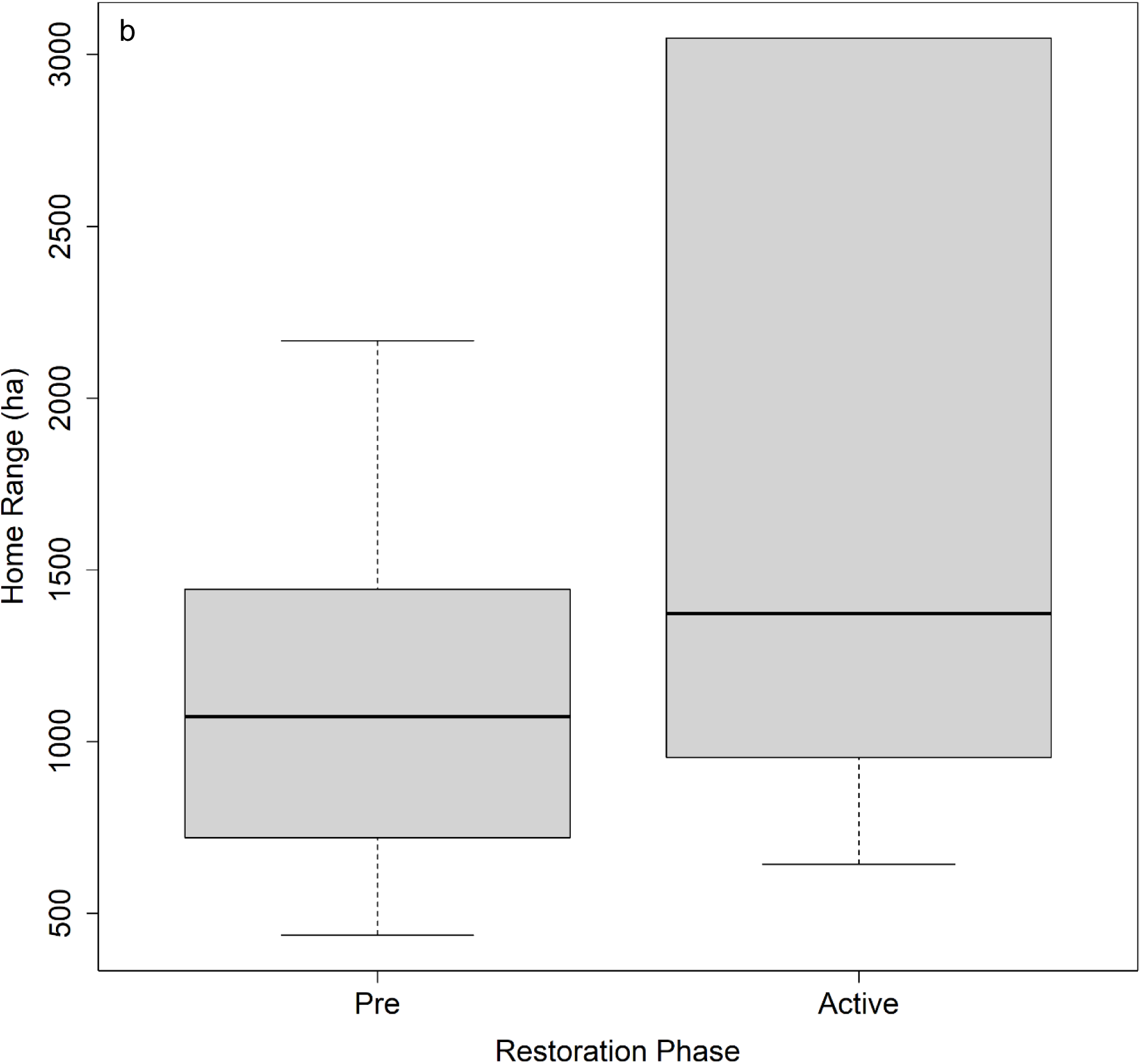
Home range size (ha) of Piping Plovers pre-restoration and during restoration (active) at Caminada Headland, Louisiana based on kernel density estimates. Displayed are the (a) 50% isopleths and (b) 95% isopleths. Central black line indicates median, top and bottom of box indicate interquartile range, and whiskers indicate total range. Due to small samples sizes during the restoration (active) phase, the maximum range and upper quartile are the same. We found no significant differences between restoration phases.

The Piping Plover resight data from Whiskey Island were limiting, such that we were only able to compare MCPs for the pre-restoration and active restoration phases (Table 1; Figures S5-S6). The lack of resights during the post-restoration phase is surprising as we observed more Piping Plovers (banded and unbanded combined) during post-restoration (n = 443) compared to active restoration (n = 384). When analyzing the 50% isopleths, home range area was similar between the pre-restoration (3.57 ± 6.84 ha) and active restoration (3.60 ± 5.71 ha) phases (χ^2^ = 0.19; df = 1; *p* = 0.66; Figures 4a and S5). The 95% isopleth exhibited a similar trend except the pre-restoration phase (12.15 ± 17.25 ha) showed less variation compared to active restoration (12.87 ± 18.60 ha; χ^2^ = 0.27; df = 1; *p* = 0.60; Figures 4b and S6).

**Figure 4.**
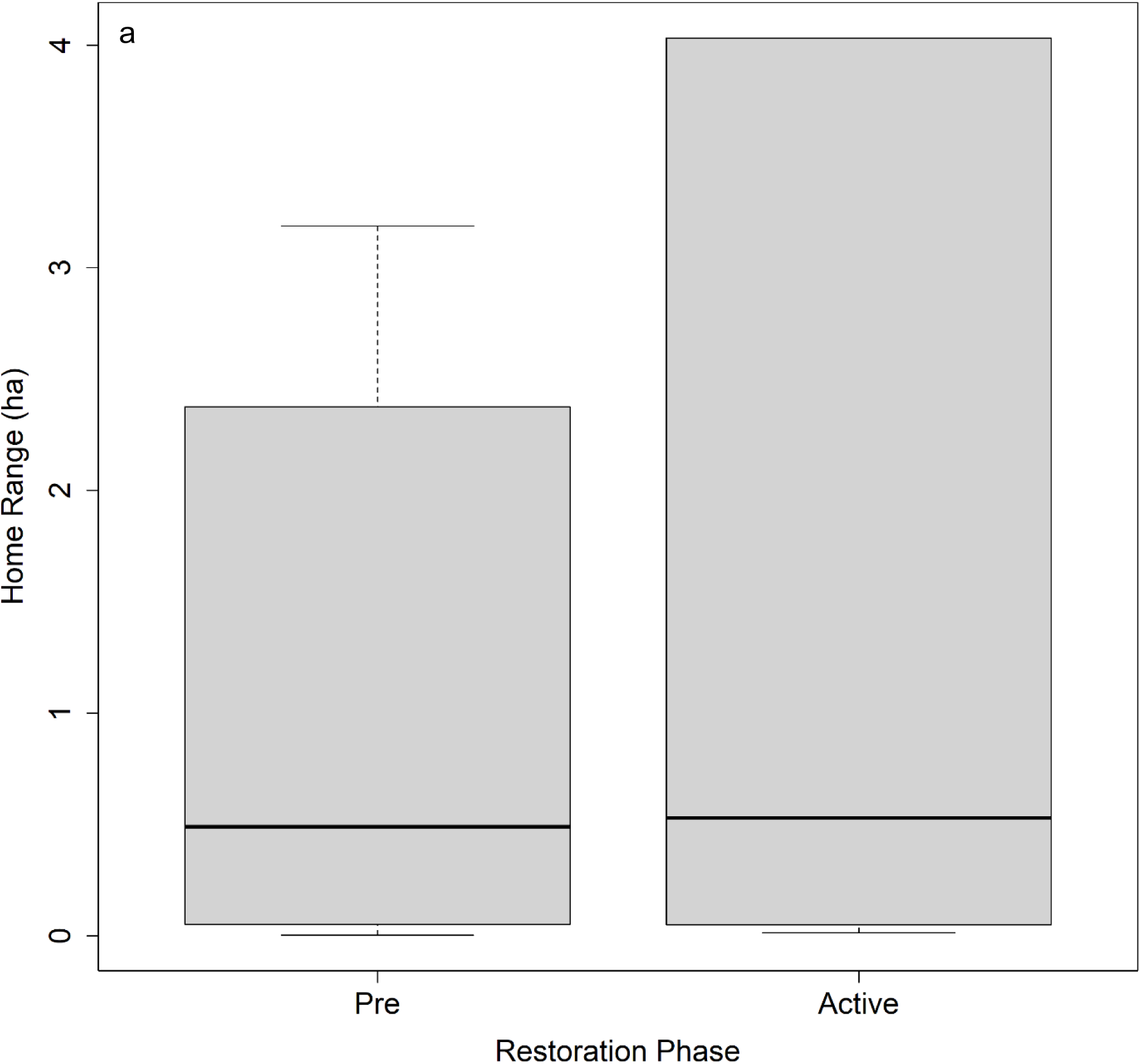
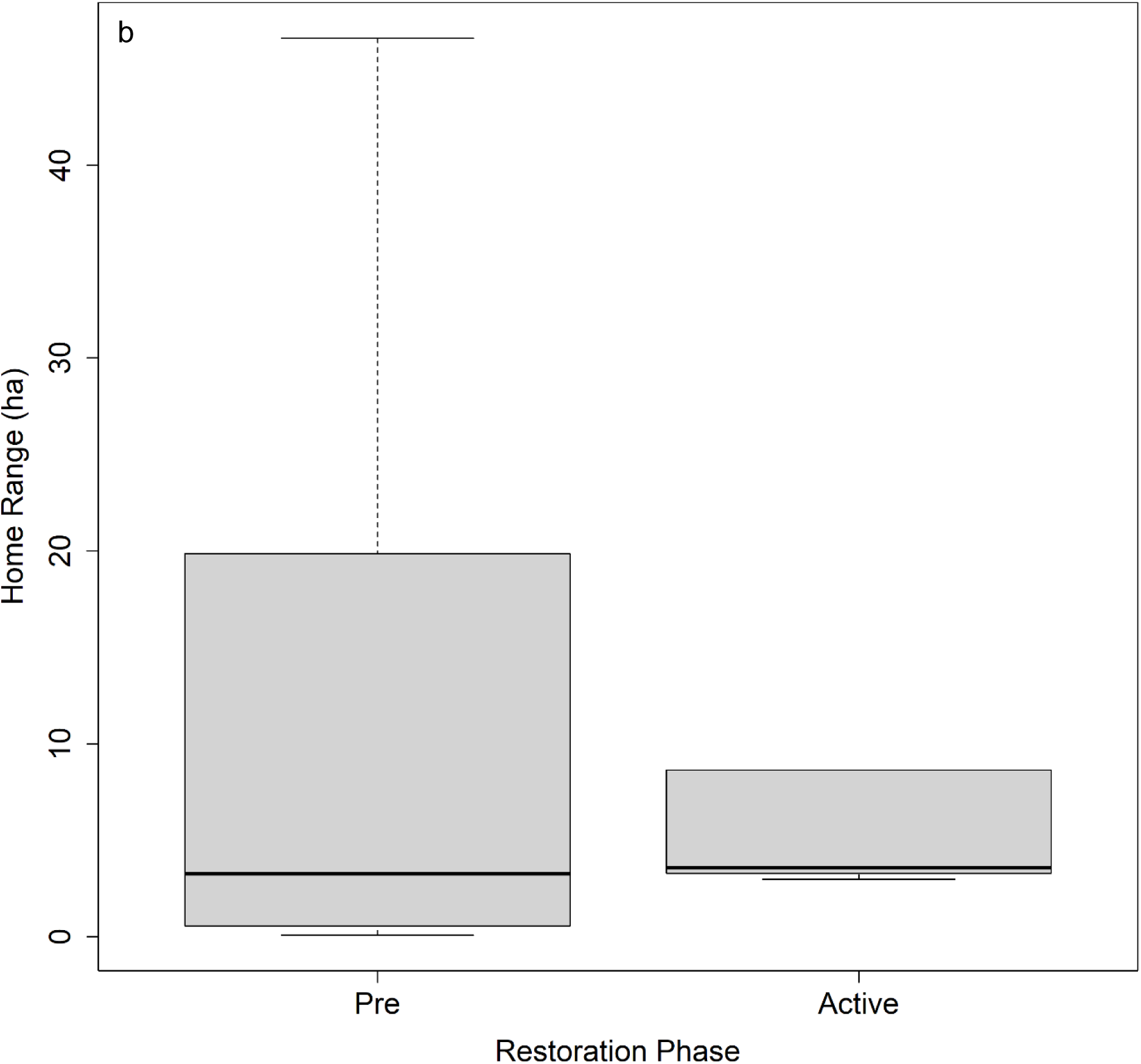
Home range size (ha) of Piping Plovers pre-restoration and during restoration (active) at Whiskey Island, Louisiana based on minimum convex polygon. Displayed are the (a) 50% isopleths and (b) 95% isopleths. Central black line indicates median, top and bottom of box indicate interquartile range, and whiskers indicate total range. Due to small samples sizes during the restoration (active) phase, the maximum range and upper quartile are the same. We found no significant differences between restoration phases.

## Discussion

Understanding spatiotemporal changes in non-breeding habitat use following restoration activities is important as such projects become more urgent and widespread. While we found little change in Piping Plover home range size among restoration phases, it is notable that individuals at Caminada Headland appear to have larger MCP-derived home ranges than those at Whiskey Island. The weak response of birds to restoration activities shows that birds are using similar or smaller amounts of habitat after restoration is complete. This result leads to two mutually exclusive hypotheses: Piping Plovers did not need to expand their foraging range following restoration, or Piping Plovers were unable to expand their home ranges. The inability of Piping Plovers to expand their home ranges could be due to intraspecific or interspecific competition due to limited resources. However, further study into differences in intertidal habitat availability, benthic invertebrate recovery, and energetic condition before and after restoration can further elucidate short-term restoration effects on site use, prey availability, and foraging as home range size alone might not be sufficient to capture changes from restoration activities.

At Caminada Headland, we had enough data to compare all three restoration phases using the MCP approach, but were only able to compare the pre- and during restoration phases using the KDE approach due to data limitations. We only found a significant change when using the MCP approach to evaluate core habitat use (50% isopleth), which showed a reduced home range size after restoration compared to before restoration occurred. The reduction in home range size following restoration may suggest that Piping Plovers are able to meet their needs within a smaller core spatial area following restoration. While we were unable to find assessments of Piping Plover home range relative to coastal restoration efforts, previous studies have found that habitat restoration projects tended to benefit Piping Plover foraging along the Atlantic Coast (Maslo et al. 2012; McIntyre and Heath 2011; Schupp et al. 2013; Smith et al. 2020). However, it should be noted that the restoration projects we cite occurring on the Atlantic coast aimed to improve Piping Plover habitat, which differed from the objectives of the barrier island and headland restoration occurring as part of this study. It is possible that foraging habitat improved for Piping Plovers at Caminada Headland following restoration, conceivably leading to smaller core home ranges, but further investigation of foraging success would be needed to test this hypothesis.

Alternatively, competition over limited resources within or among species could also limit home range size, but again more information is needed to determine recovery of benthic organisms and degree of competition to support or refute these hypotheses. The home ranges observed at Caminada Headland compare variably to published results (Cohen et al. 2008; Drake et al. 2001; Noel and Chandler 2008) depending on the approach. Our analysis with MCP found home ranges to be smaller than published values, while the KDE approach estimated home ranges to be larger than published values. These differences may be due to analytical or field methods used, but could also reflect biologically meaningful differences between Piping Plovers on the Atlantic Coast versus the Gulf. For example, factors such as prey abundance, foraging substrate, anthropogenic use, competition, or weather could conceivably influence home range size between different geographic locations.

At Whiskey Island, we were restricted to comparing only the pre- and active phases of restoration and did not find any difference in home range sizes. Home range sizes on Whiskey Island were substantially smaller than other published home range sizes regardless of restoration phase (Cohen et al. 2008; Drake et al. 2001; Noel and Chandler 2008), but this difference may be due to non-biological reasons, such as lack of anthropogenic disturbance, methods used to calculate home range sizes, limitation of our study to one island within a barrier island chain, amount of habitat, or lack of a banding program at Whiskey Island during survey years. While our survey efforts were restricted to Whiskey Island, future research might focus on a comprehensive survey of the Isle Dernieres Barrier Islands Refuge to determine the spatial distribution and movement ecology of Piping Plovers across the entire barrier island chain. Moreover, additional monitoring of Whiskey Island could generate larger sample sizes to improve comparisons between pre- and post-restoration phases.

There were multiple Piping Plovers that were observed during multiple seasons, which provides some additional information on individual responses to the restoration activities taking place at our study sites (e.g., Figures S7-S9). While this was more limited at Whiskey Island with only one individual being detected during more than one restoration phase (Figure S9), there were a higher number of repeat observations at Caminada Headland when considering MCP derived home ranges. Individuals tended to use similar geographic areas across seasons and restoration phases (Figures S7-S9). However, home range size within (i.e., same phase, different seasons) and among restoration phases showed substantial variability. There were also differences based on the isopleth size (i.e., 50% or 95%), which impacted the direction of the relationship of home range size among restoration phases. Anecdotally, the high within-individual variability in home range size further suggests that home range size alone is likely insufficient at assess restoration impacts on Piping Plovers and additional metrics could help in assessing restoration impacts.

To our knowledge, this is the first study to determine if and how Piping Plover non-breeding home ranges change following habitat restoration. Our results suggest that barrier island and headland restoration along the Louisiana coast did not meaningfully impact Piping Plover habitat use, which is valuable as land managers in Louisiana and across the Gulf continue to restore and create coastal habitats (Brush et al. 2019; Coastal Protection and Restoration Authority of Louisiana 2023; Texas General Land Office 2023). However, we do acknowledge that this research has limitations due to small sample sizes that did not allow us to compare among all restoration phases, necessitating the need for a more robust study that can compare home range size across all restoration phases. Moreover, being able to compare sites before and after restoration with “control” sites over that same temporal period would be particularly powerful, although finding appropriate control sites may be difficult considering the dynamic geomorphology of coastal habitats on relatively short temporal scales. Given the amount of restoration slated for areas of the Gulf and beyond (e.g., Coastal Protection and Restoration Authority of Louisiana 2023; Deepwater Horizon Natural Resource Damage Assessment Trustees 2016; Deepwater Horizon Oil Spill Regionwide Trustee Implementation Group 2021; Texas General Land Office 2023), it is important to have an understanding of how restoration practices can impact species and how to design restoration projects that can meet objectives to protect human life and infrastructure while also providing suitable habitat for species of conservation concern. Our ability to work at a headland and barrier island allowed us to investigate restoration activities at different geographic features with slightly differing restoration methodology. Additional studies on habitat succession, foraging success, and physiology could further verify that Piping Plovers are meeting energetic needs and not falling into an ecological trap at restoration sites, given that Piping Plovers using disturbed sites tend to have lower energetic condition and survival compared to birds using less disturbed sites (Gibson et al. 2018).

## Acknowledgements

We appreciate helpful comments from D. Lee, J. Sylvest, P. Leberg, N. Enwright, and anonymous reviewers on earlier drafts of this manuscript. We also appreciate J. Smolinsky, N. Enwright, C. Kingwill, H. Thurman, and L.A. Randall for assistance with figures; J. Smolinsky and D. Johnson for their support in developing R code; W.C. Barrow and C. Jeske for assisting with project management; L.A. Randall for assistance with the data release; and J. Schulz as well as volunteers for assisting with data collection. We thank the groups banding Piping Plover used in this study, which includes The Tern and Plover Conservation Partnership; Nebraska Game and Parks Commission; Virginia Tech Shorebird Program; U.S. Geological Survey Northern Prairie Wildlife Research Center; New Jersey DEP Fish and Wildlife Endangered and Nongame Species Program; Environment and Climate Change Canada; and State University of New York, College of Environmental Sciences and Forestry. Information about Piping Plovers banded in the Great Lakes were obtained from the Great Lakes Piping Plover database maintained at the University of Minnesota-Twin Cities. This work was supported by the CPRA (contract number: 2000387259), the Barataria-Terrebonne National Estuary Program, and the U. S. Geological Survey. Any use of trade, firm, or product names is for descriptive purposes only and does not imply endorsement by the U. S. Government.

## Notes

https://doi.org/10.5066/P9LQ7JKZ

### Competing Interest Statement

The authors have declared no competing interest.

