## Supplemental Information for "Piping Plover Home Ranges Do Not Appear to be Impacted by Restoration of Barrier Islands and Headlands"

*This information product has been peer reviewed and approved for publication as a preprint by the U.S. Geological Survey.*

### Supplemental Information for: Piping Plover Home Ranges do Not Appear to be Impacted by Coastal Restoration of Barrier Islands and Headlands

Theodore J. Zenzal Jr. [1*], Amanda N. Anderson[1§], Delaina LeBlanc[2], Robert C. Dobbs[1‡], Brock Geary[1†], and J. Hardin Waddle[3]

[1]U.S. Geological Survey, Wetland and Aquatic Research Center, Lafayette, LA

[2]Barataria-Terrebonne National Estuary Program, Thibodaux, LA

[3]U.S. Geological Survey, Wetland and Aquatic Research Center, Gainesville, FL

[§]Present address: Gulf South Research Corporation, Baton Rouge, LA

[‡]Present address: Louisiana Department of Wildlife and Fisheries, Lafayette, LA

[†]Present address: Department of Pathobiology, Wildlife Futures Program, University of Pennsylvania School of Veterinary Medicine, Kennett Square, PA

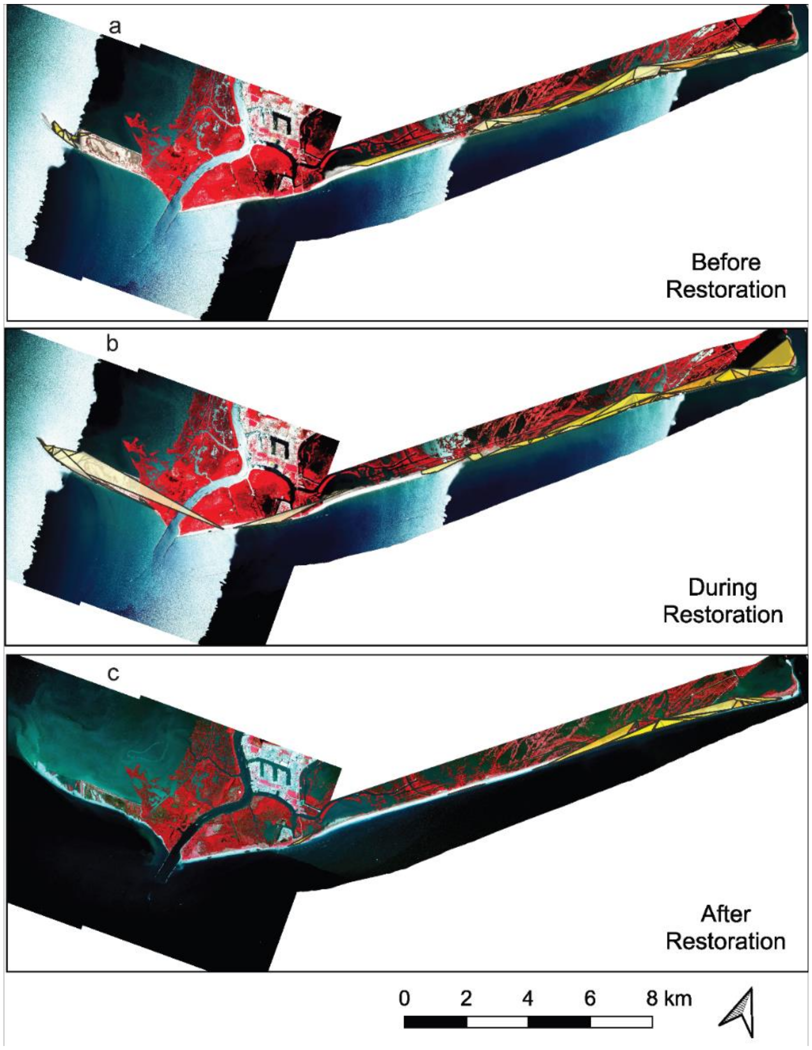


Figure S1: Seasonal Piping Plover home ranges based on the minimum convex polygon using a 50% isopleth (a) before restoration, (b) during restoration, and (c) after restoration at Caminada Headland. Imagery is shown as false color, which means redder areas indicate greener vegetation. Some individuals were detected multiple years, thus contributing more than one home range when data were available to produce a home range for each year they were observed (refer to article for details). Basemap provided by the USDA National Agriculture Imagery Program.

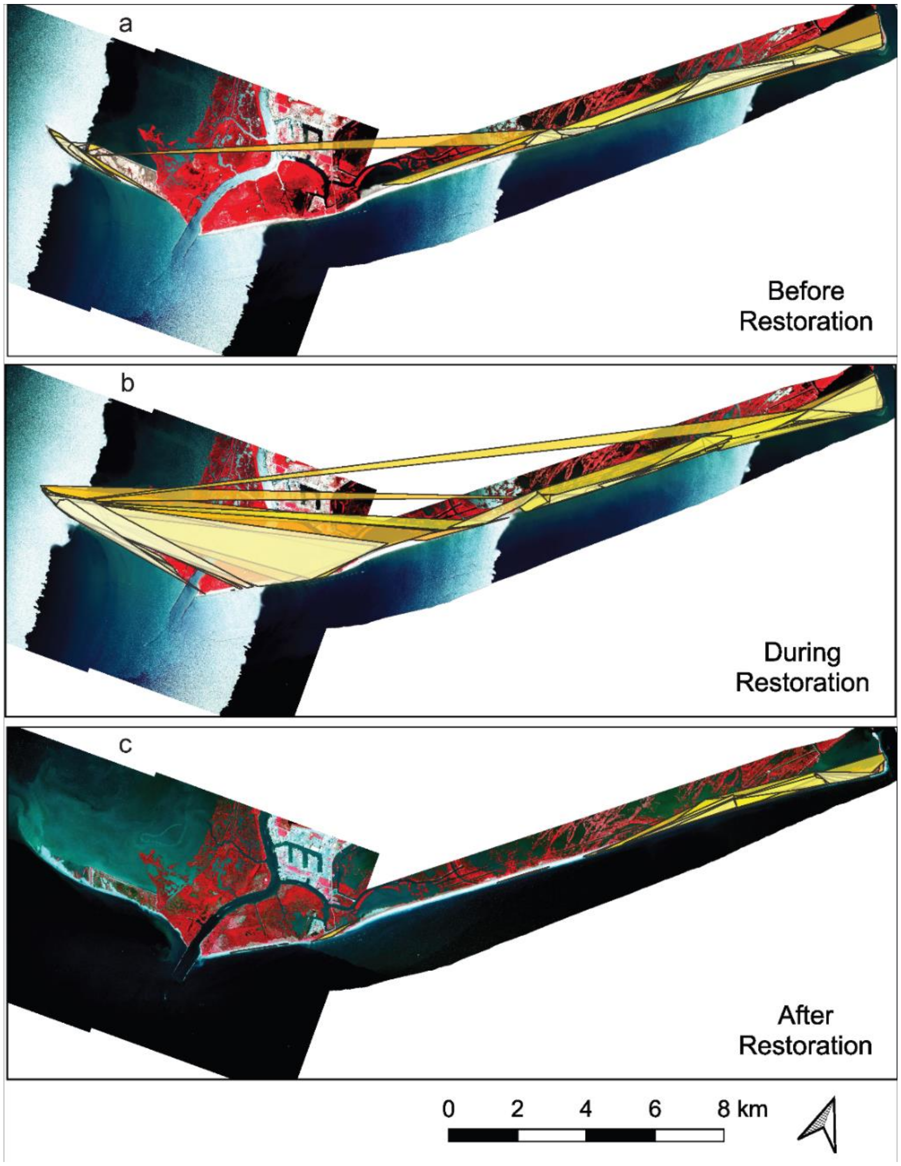


Figure S2: Seasonal Piping Plover home ranges based on the minimum convex polygon using a 95% isopleth (a) before restoration, (b) during restoration, and (c) after restoration at Caminada Headland. Imagery is shown as false color, which means redder areas indicate greener vegetation. Some individuals were detected multiple years, thus contributing more than one home range when data were available to produce a home range for each year they were observed (refer to article for details). Basemap provided by the USDA National Agriculture Imagery Program.

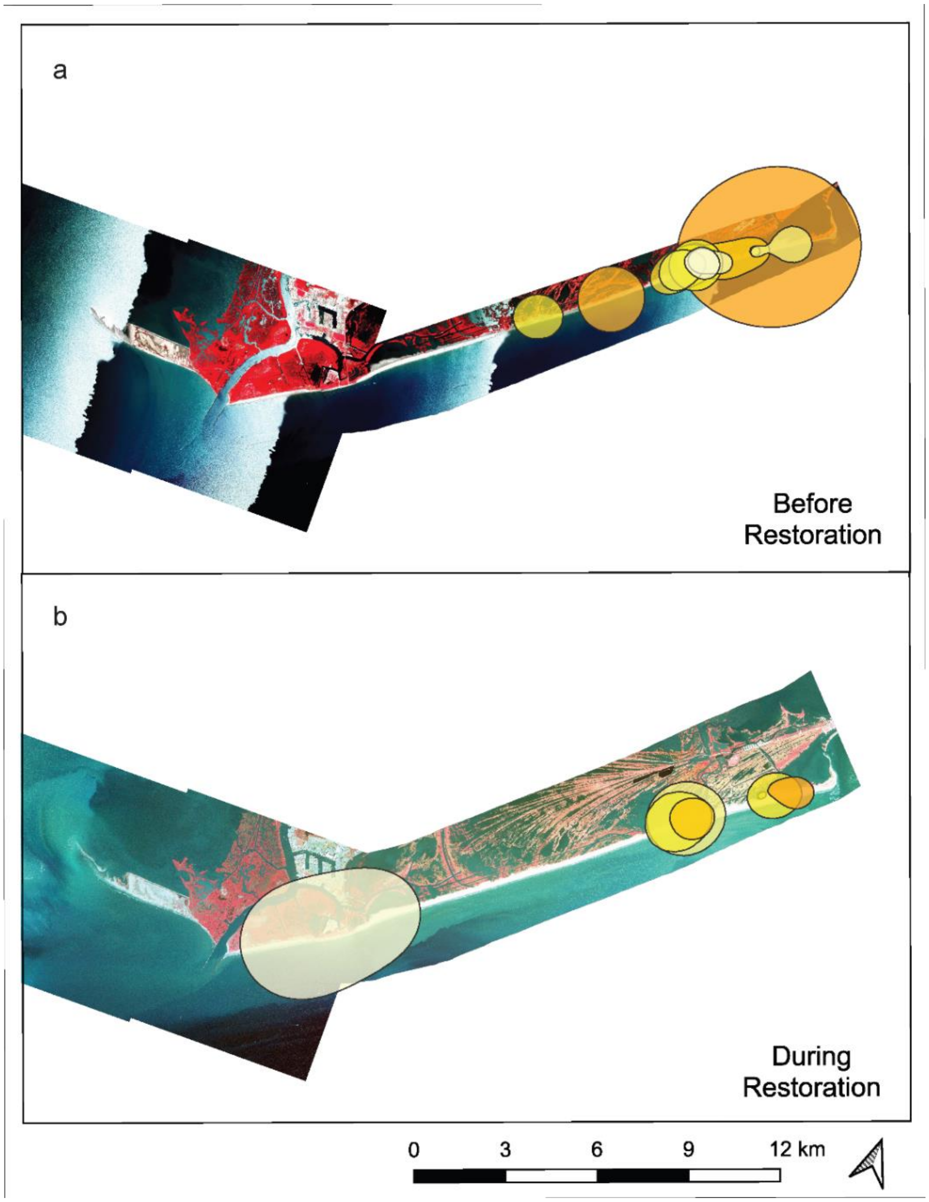


Figure S3: Seasonal Piping Plover home ranges based on the kernel density estimate using a 50% isopleth (a) before restoration and (b) during restoration at Caminada Headland. Imagery is shown as false color, which means redder areas indicate greener vegetation. Some individuals were detected multiple years, thus contributing more than one home range when data were available to produce a home range for each year they were observed (refer to article for details). Basemap provided by the USDA National Agriculture Imagery Program.

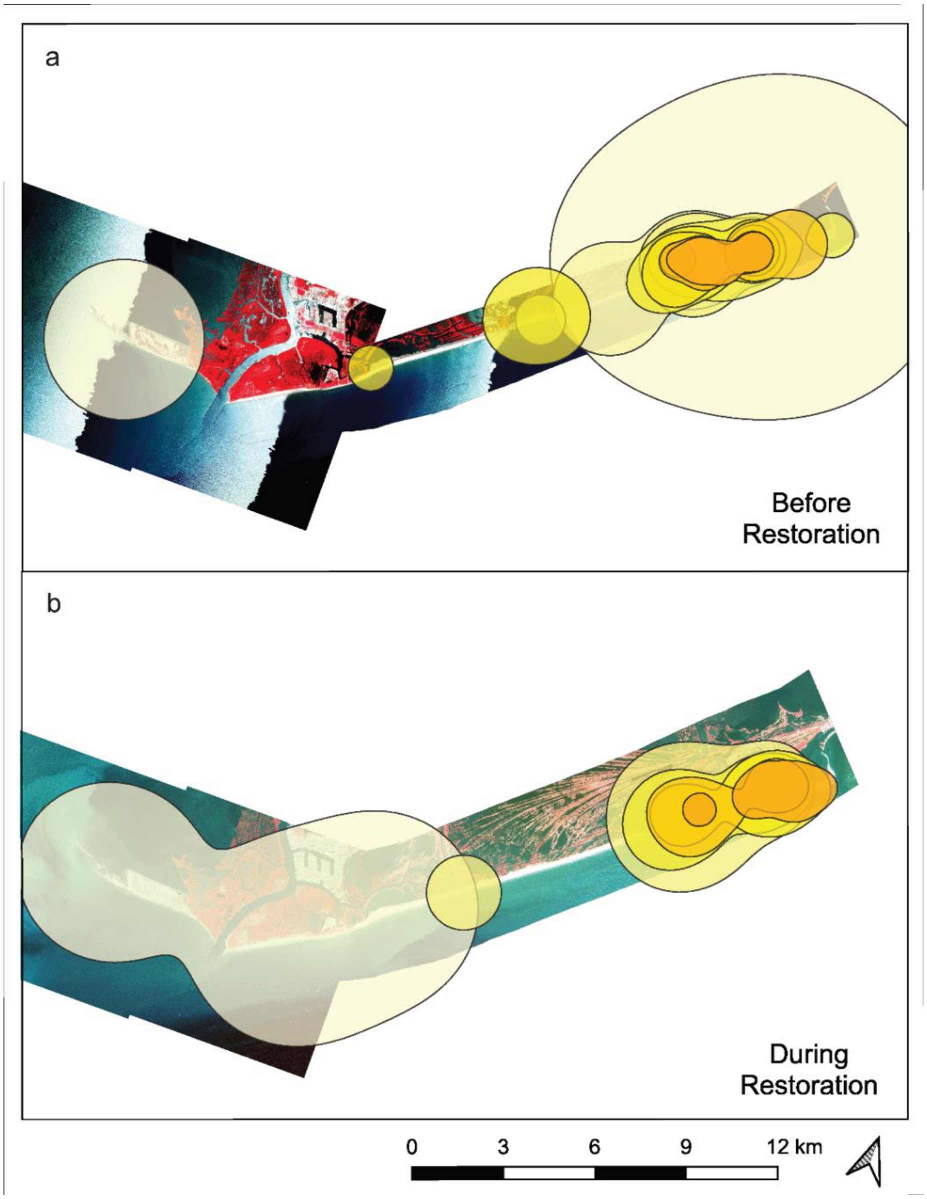


Figure S4: Seasonal Piping Plover home ranges based on the kernel density estimate using a 95% isopleth (a) before restoration and (b) during restoration at Caminada Headland. Imagery is shown as false color, which means redder areas indicate greener vegetation. Some individuals were detected multiple years, thus contributing more than one home range when data were available to produce a home range for each year they were observed (refer to article for details). Basemap provided by the USDA National Agriculture Imagery Program.

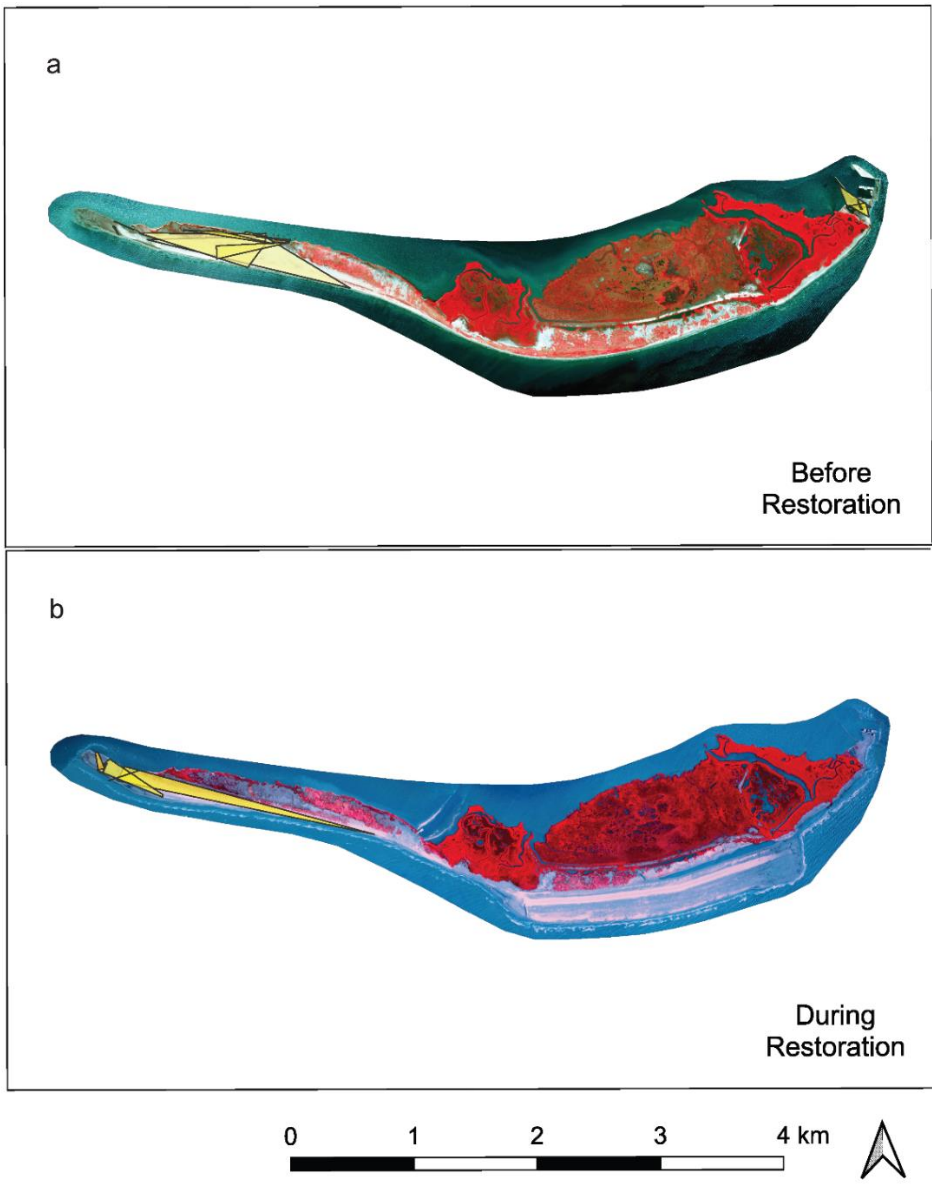


Figure S5: Seasonal Piping Plover home ranges based on the minimum convex polygon using a 50% isopleth (a) before restoration and (b) during restoration at Whiskey Island. Imagery is shown as false color, which means redder areas indicate greener vegetation. Some individuals were detected multiple years, thus contributing more than one home range when data were available to produce a home range for each year they were observed (refer to article for details). Basemap provided by the USDA National Agriculture Imagery Program.

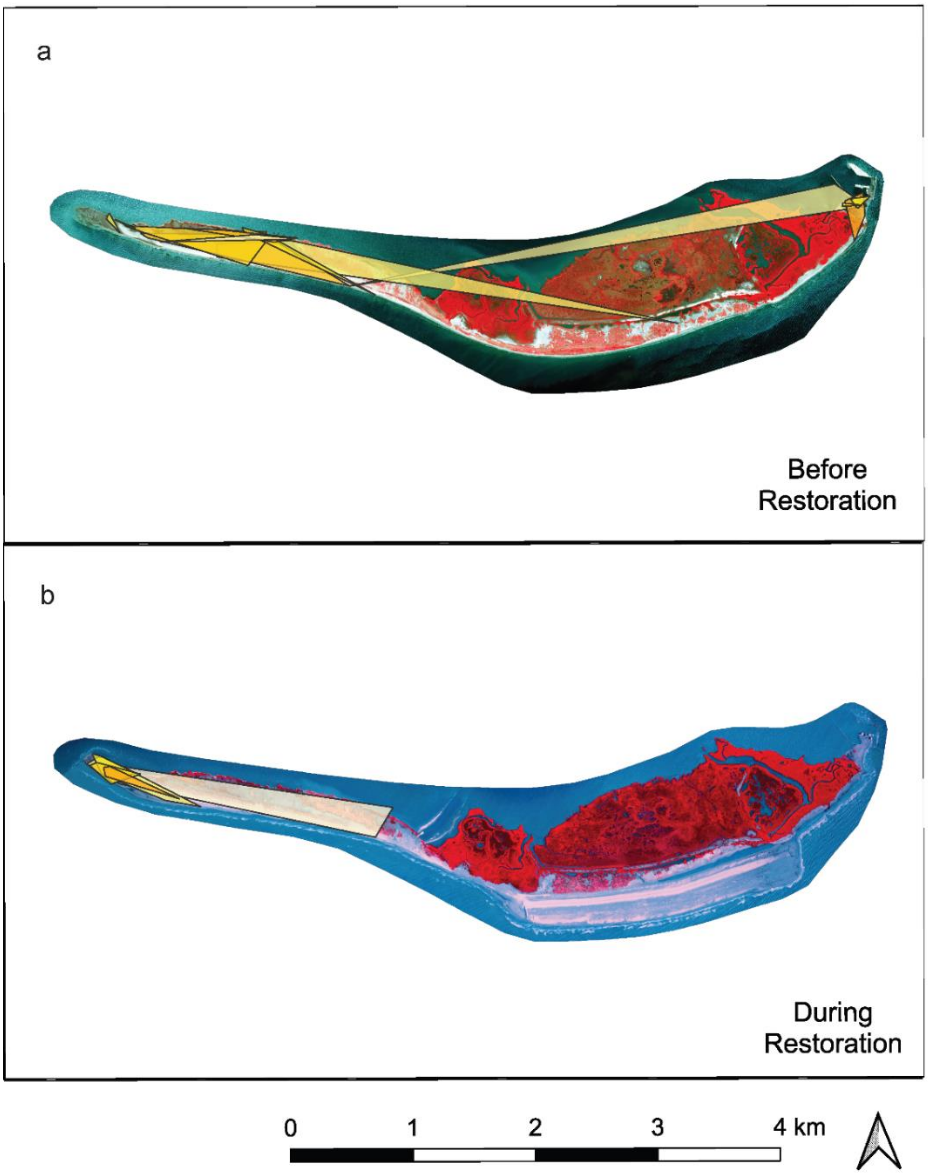


Figure S6: Seasonal Piping Plover home ranges based on the minimum convex polygon using a 95% isopleth (a) before restoration and (b) during restoration at Whiskey Island. Imagery is shown as false color, which means redder areas indicate greener vegetation. Some individuals were detected multiple years, thus contributing more than one home range when data were available to produce a home range for each year they were observed (refer to article for details). Basemap provided by the USDA National Agriculture Imagery Program.

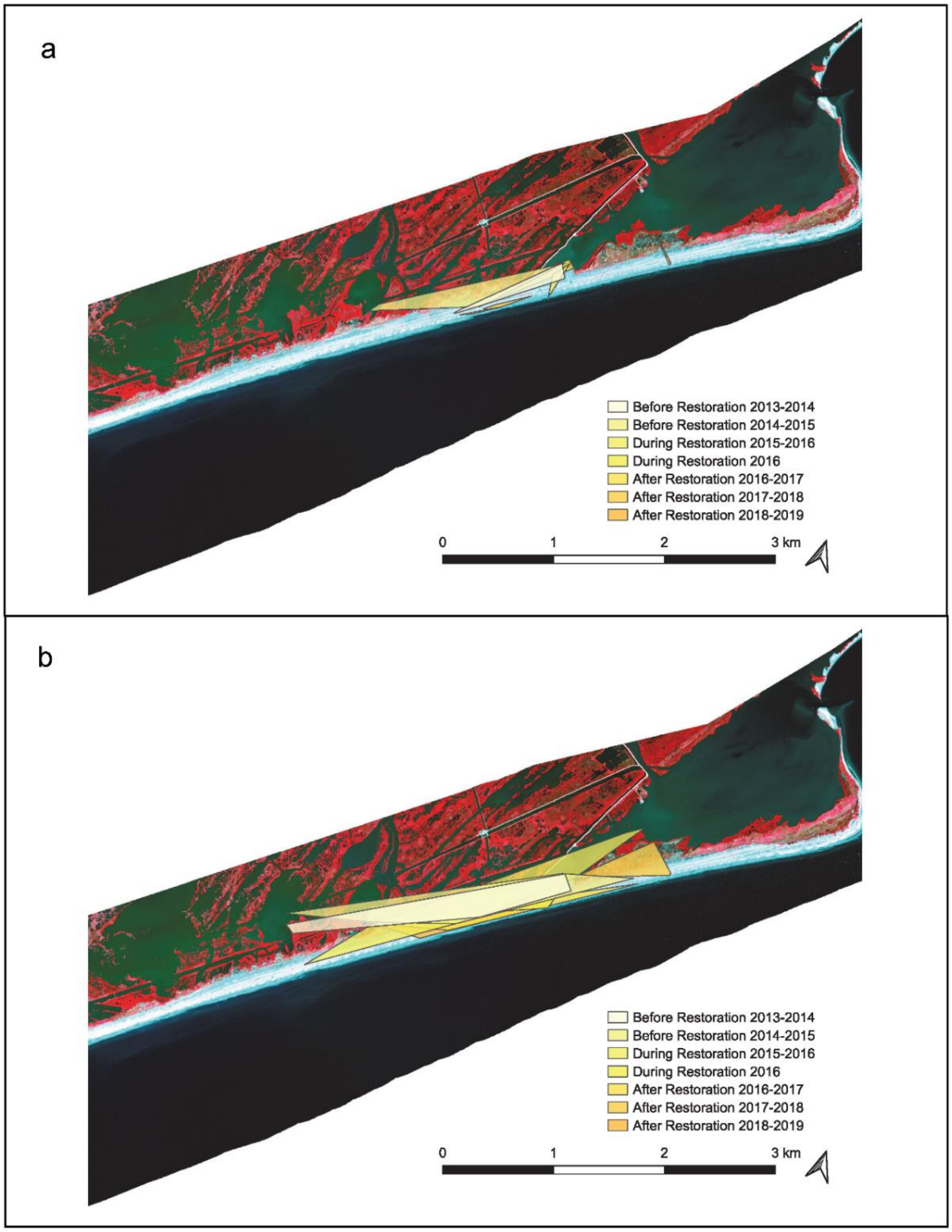


Figure S7: Home range of an individual Piping Plover based on the minimum convex polygon before and during restoration at Caminada Headland. The (a) 50% and (b) 95% isopleths are shown as false color, which means redder areas indicate greener vegetation. Basemap provided by the USDA National Agriculture Imagery Program.

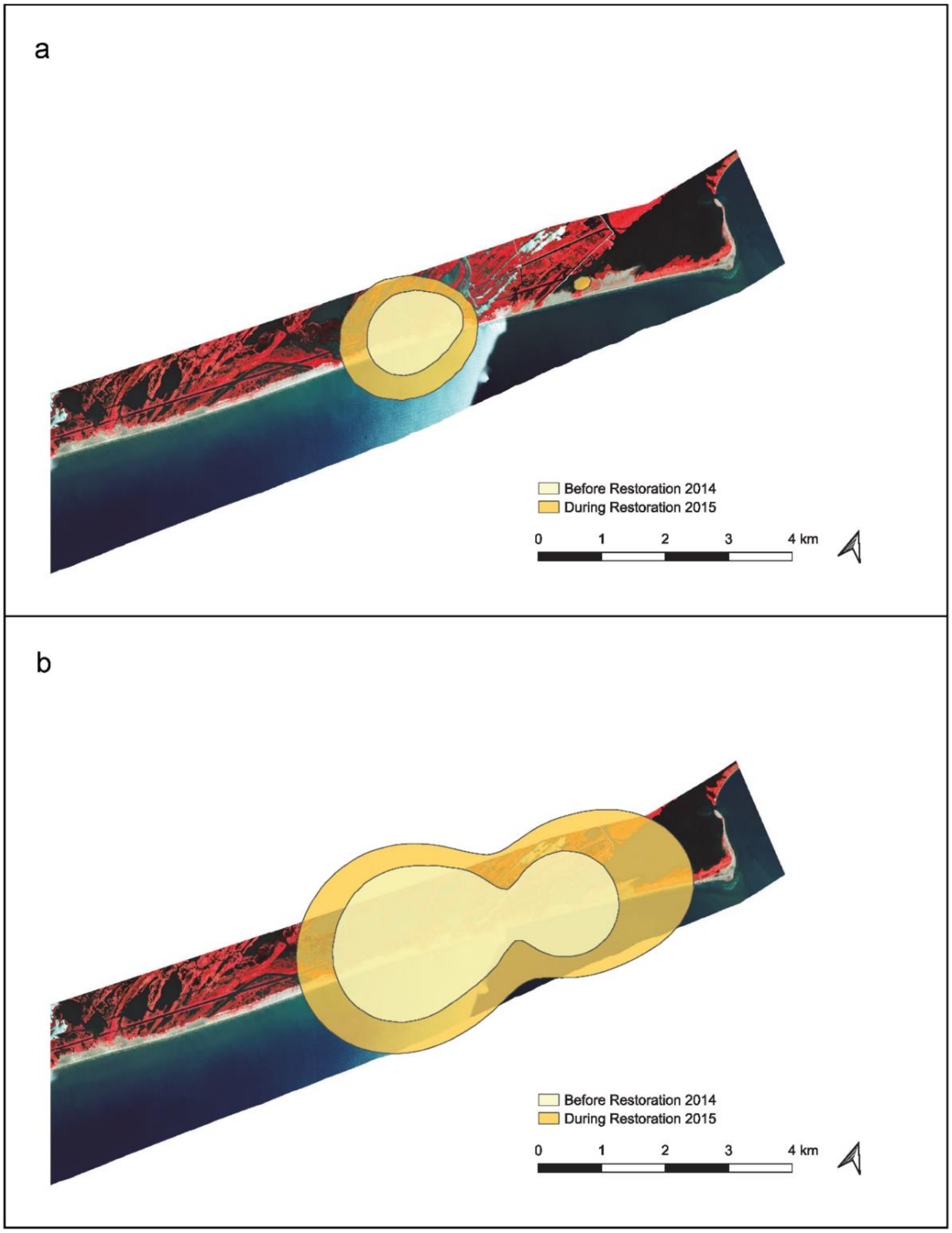


Figure S8: Home range of an individual Piping Plover based on the kernel density estimate before and during restoration at Caminada Headland. The (a) 50% and (b) 95% isopleths are shown as false color, which means redder areas indicate greener vegetation. Basemap provided by the USDA National Agriculture Imagery Program.

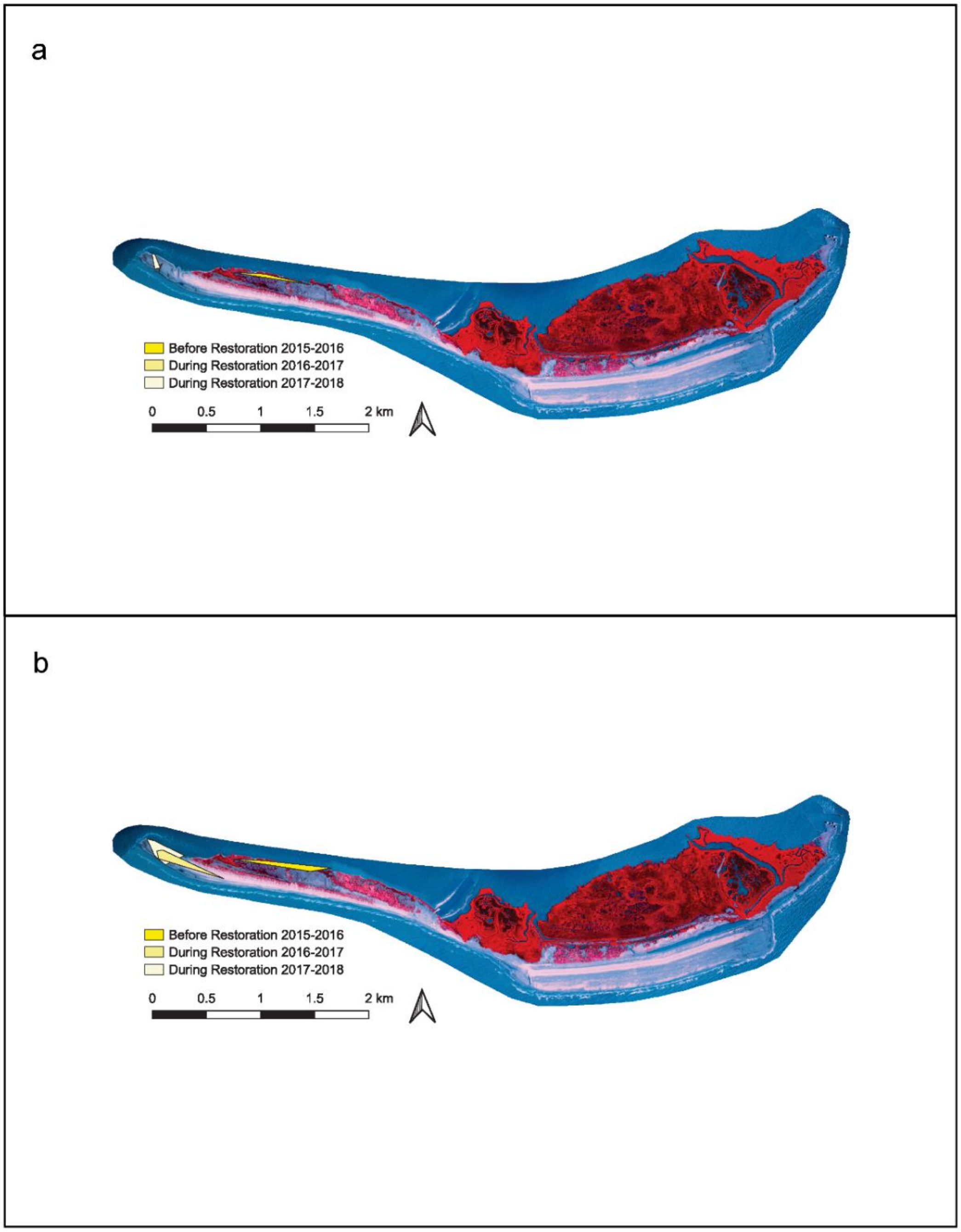


Figure S9: Home range of an individual Piping Plover based on the minimum convex polygon before and during restoration at Whiskey Island. The (a) 50% and (b) 95% isopleths are shown as false color, which means redder areas indicate greener vegetation. Basemap provided by the USDA National Agriculture Imagery Program.
